# Environmental history can counteract nutrient quality in shaping the evolution of bacterial growth rates

**DOI:** 10.64898/2026.09.17.752447

**Authors:** Arjun Mandyam Dhati, Jacopo Grilli, Akshit Goyal

## Abstract

Despite sharing nearly identical enzymes, strains of the same microbial species display wide variability of growth rates when growing on the same nutrients. Here, we show that such growth rate variability can emerge as a consequence of evolutionary adaptation to different patterns of environmental fluctuations. We develop a mathematical model combining cellular proteome allocation with eco-evolutionary dynamics in fluctuating environments We find that different patterns of environmental fluctuations select for distinct growth profiles. Slow-growing strains repeatedly outcompete fast-growers when evolved in rapidly changing environments. In a range of environments with asymmetric nutrient availability, evolution reproducibly leads to the robust diversification of two coexisting strains. We develop a theoretical framework based on adaptive dynamics which quantitatively predicts these evolutionary outcomes and explains them in terms of the growth-lag tradeoff that all cells face. Finally, we show that the model can reproduce several patterns in growth-rate data from closely-related strains. Our results highlight that bacterial growth rates can rapidly adapt to their recent environmental history by proteome allocation alone, in ways that often oppose expectations based on nutrient quality.

## Introduction

Bacteria in nature vary widely in their growth rates [1, 2, 3]. Indeed, growth rate differences often form the basis of how bacteria are classified, from slow-growing oligotrophs to fast-growing copiotrophs [4]. To explain the origin and maintenance of this growth rate variability, two factors are typically invoked. The first is nutrient quality: richer nutrients support higher metabolic fluxes and energetic yields, allowing faster growth [5, 6, 7]. The second is enzymatic machinery: some species may genetically encode enzymes that are faster or more efficient [8, 9]. Together, these factors have traditionally been used to explain how growth rates differ across taxa and environments.

However, neither nutrient quality differences nor enzymatic differences can explain growth variability between strains of the same bacterial species can exhibit substantial variation in growth rates even on the same nutrient [3, 10, 11, 12, 13]. For instance, Fig. 1a shows the growth rates of strains of *E. coli* and *V. splendidus* on two different nutrients. Not only do we observe substantial growth rate variability, we also observe that different strains can show inverted patterns across nutrients. In Fig. 1a, one strain of *V. splendidus* (marked in red) is the slowest grower on glucose but the fastest on pyruvate, while another strain (also marked in red) shows the opposite: it is the fastest grower on glucose but slowest on pyruvate (also see [3]). These observations cannot be attributed to enzymatic differences alone, as such strains can differ only by a few mutations in some cases, as well as code for the same metabolic enzymes [14, 15, 16, 17]. Thus, at the strain level, we are left with a puzzle: how can growth rate variability rapidly emerge in a way that appears to counteract nutrient quality and enzymatic differences?

**Figure 1.**
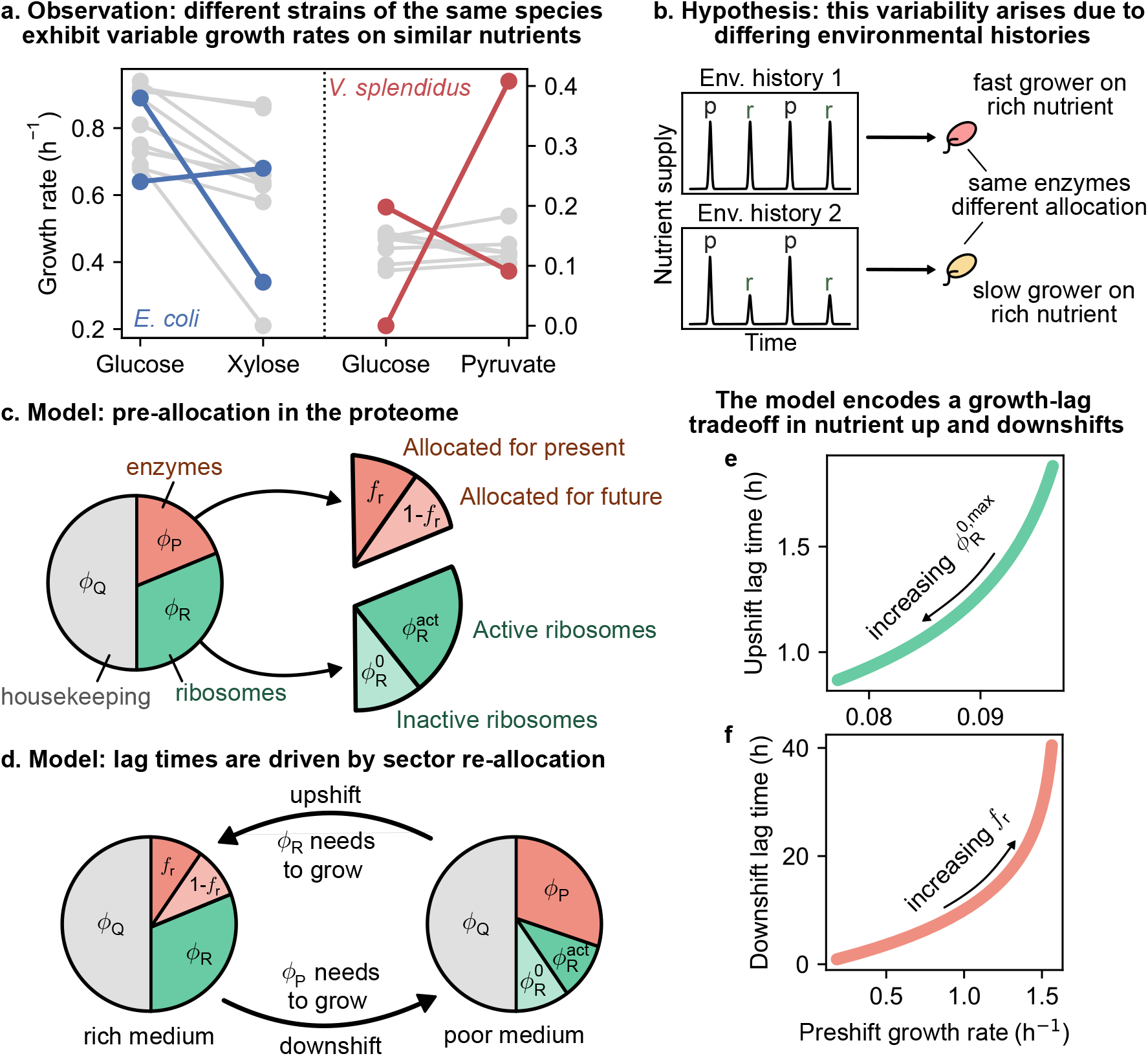
Growth rate variability across strains motivates a model of proteome allocation in fluctuating environments. **a** Different strains of *E. coli* and *V. splendidus* differ in their growth rates on the same nutrients, suggesting growth rate is a property of the strain rather than the nutrient. Highlighted are two strains for each species that show a contrasting trend in growth rates across nutrients. Data for *E. coli* from Ref. [11], *V. splendidus* from [3]; see SI. **b** We hypothesize that this behaviour emerges from differing evolutionary histories of the strains. **c** The bacterial proteome can be divided into three sectors, namely P (metabolic enzymes), R (ribosomes) and Q (housekeeping). Cells may pre-allocate for future conditions in both the P and R sectors. **d** Optimal allocation dictates a large R sector during fast growth and a large P sector during slow growth. A lag phase during nutrient shifts arises from the need to re-allocate the proteome upon encountering a new nutrient. **e, f** Our model captures the growth-lag tradeoff for both nutrient upshifts and downshifts.

A natural but underexplored explanation for this variability is that growth rates do not reflect adaptation to a fixed condition, but they rather reflect a compromise between optimizing growth in the present condition and preparedness for future uncertain environments [18, 19, 20, 21, 22]. In fact, in fluctuating environments, maximizing growth rate in a given condition might come at a cost when that condition changes [23, 24, 25, 26]. As illustrated in Fig. 1b, two strains with identical metabolic capabilities can nonetheless grow at different rates on the same nutrient if they differ in how they allocate internal cellular machinery based on past environmental exposure [27, 28]. A population frequently exposed to a given condition may rapidly evolve its allocation to exploit this condition, whereas another strain population, exposed to a different sequence of environments, may evolve to allocate its cellular resources differently in preparedness for the changes in the environment [6, 29, 30]. In this view, growth rates are not solely determined by the present environment, but can rapidly evolve to reflect the environments it has been exposed to over evolutionary time [24]. Despite this intuitive hypothesis, the role of environmental history in shaping growth rate variability remains poorly understood [21, 26]. In particular, there is no predictive framework that quantitatively links environmental fluctuations, evolution of cellular allocation strategies, and the resulting growth rates of strains.

Here, we develop such a framework to predict growth rate variability from the environmental history that strains evolved in. Our framework combines proteome allocation theory [5, 6, 31, 32] with fluctuating environments and eco-evolutionary dynamics [33, 34, 35, 36, 37]. Specifically, in contrast with previous work focusing on proteomic changes during nutrient upshifts [31] or between equivalent nutrients [37], we allow populations to pre-allocate the proteome in anticipation of both nutrient upshifts and downshifts, capturing the full spectrum of environmental change. We show that distinct patterns of environmental fluctuations select for distinct growth-rate profiles, even among strains with identical biochemical capabilities and on resources with the same quality. Our model recapitulates the observed growth-rate variability in the data (Fig. 5b), and makes two testable predictions: (1) slow-growing strains are favored in environments with higher relative supply of poor nutrients, (2) strain coexistence is favored in environments with asymmetric rich and poor nutrient availability. Our results provide a quantitative link between environmental history and microbial growth, suggesting that growth rate profiles observed in natural populations reflect environmental fluctuation patterns rather than purely biochemical constraints.

## Results

### A model of proteome allocation in fluctuating environments

We use a coarse-grained description of the bacterial proteome to model the dependence of growth rates on environmental conditions. Following proteome allocation theory, we divide the proteome into three sectors: a metabolic sector *ϕ*_P_ that imports nutrients and produces precursors, a ribosomal sector *ϕ*_R_ that synthesizes protein, and a housekeeping sector *ϕ*_Q_ that captures all remaining functions, with *ϕ*_P_ + *ϕ*_R_ + *ϕ*_Q_ = 1 (Fig. 1c). At steady state, the growth rate is set by the balance between metabolic flux and protein synthesis. This balance links proteome allocation directly to growth rates and forms the basis of our framework (Methods).

To model fluctuating environments, we consider cells that repeatedly shifted between nutrient conditions. We will start with two nutrients: a higher quality rich nutrient and a lower quality poor nutrient, though we will later show that our model and results generalize to more complex environments with multiple nutrients. Following each upshift (poor to rich) or downshift (rich to poor), cells must reallocate their proteome before reaching a new steady-state growth rate. We represent this period of transient adjustment by a lag time: the time quantifying the amount of growth lost during reallocation (Fig. 1d). We parameterize allocation using two quantities: 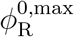, a maximal fraction of ribosomes held inactive, and *f*_r_, the fraction of metabolic enzymes dedicated to the rich nutrient when available (Methods). These parameters determine both the post-shift growth rate and the lag time by determining steady-state allocation values. For simplicity, we assume that the reallocation time of the sector that is adjusted by the largest amount sets the lag.

During an upshift, cell growth is limited by ribosomes. Cells with a larger pool of pre-allocated, inactive, ribosomes expand faster, so increasing 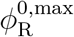 reduces lag times during up-shifts. After a downshift, cells need to synthesize enzymes related to the metabolic sector. Cells that retain enzymes for consuming the poorer nutrient reallocate faster, so decreasing *f*_r_ reduces lag times during downshifts. These same parameters also control steady-state growth rates, but in the opposite direction, generating the observed growth rate–lag tradeoff [23] (Methods). Increasing 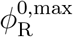 reduces the pool of active ribosomes and lowers growth rates in rich environments. Decreasing *f*_r_ reduces the investment of enzymes in the rich nutrient and also lowers the growth rates in rich media (Fig. 1e,f). As a result, no allocation strategy can simultaneously achieve maximal growth rates across nutrient conditions. Different strategies, i.e., combinations of *f*_r_ and 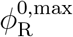, perform differently depending on the availability of rich and poor nutrients in the environment.

In this work, we focus on evolution in fluctuating environments. On these timescales, we assume that populations can change their growth rates by evolving their proteome allocation strategies rather than their metabolic enzymes efficiencies or rates. This is because regulatory changes can reshape proteome allocation and growth rates with just a few mutations even when there are no detectable changes in enzyme kinetics [38, 14], which might instead require several mutations. Motivated by these observations, throughout this work, we only allow *f*_r_ and 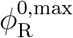 to vary over evolution, and keep all other cellular properties fixed. In this way, proteome allocation provides the mechanistic link between environmental fluctuations and growth rate variation. This framework enables us to determine how fluctuation patterns select allocation strategies, and how these strategies generate differences in growth rates across strains.

### A single allocation strategy repeatably evolves under symmetric environmental fluctuations

To determine how fluctuating environments shape allocation strategies, we simulated evolution in a serial-dilution setup that alternated between rich and poor nutrients (Fig. 2a,b). Cells grew in a medium containing one of the two nutrients for a fixed time *T* . We chose this time *T* to be long enough such that the population could typically deplete the nutrient completely and enter a non-growing state. After this, we diluted the population size by a factor *D*, and transferred it to a fresh medium containing the other nutrient. This serial dilution process generated repeated boom and bust cycles, mimicking serial dilution experiments in the lab [39, 14, 40] and feast-famine cycles in natural environments [41]. We controlled environmental fluctuations through two parameters: an effective mortality rate set by the dilution factor *D*, and the relative concentration of the rich to poor nutrient, measured in units of biomass *c*_r_*/c*_p_ (Methods). We first focused on symmetric environments with an equal supply of rich and poor nutrients, *c*_r_*/c*_p_ = 1. Note that we make the choice to measure nutrients concentrations in rescaled units of biomass for convenience (or, equivalently, we set without loss of generality the yields *Y*_r_ = *Y*_p_ = 1). In these units, when *c*_r_*/c*_p_ = 1, the population ultimately grows to the same abundance in both the rich and poor nutrient environments (Fig. 2a; see Methods for details). This is why we refer to such environmental fluctuation patterns as “symmetric”.

**Figure 2.**
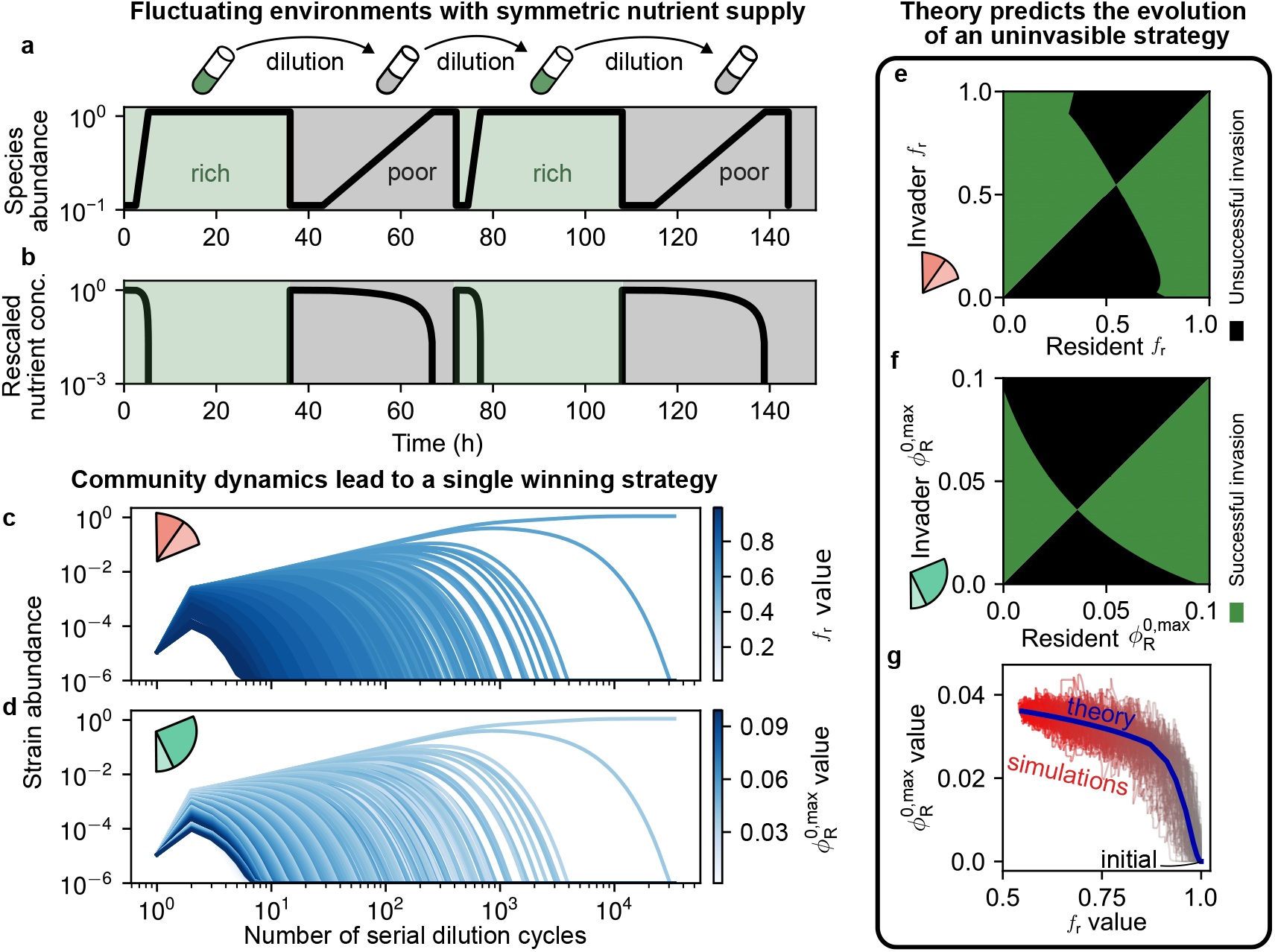
Symmetric environments select a single winning allocation strategy. **a, b** Example dynamics of a bacterial population growing in a serial dilution setup alternating between rich and poor media of equal quantities. Here, nutrient concentration ratio, *c*_r_*/c*_p_ = 1 and dilution factor *D* = 10. **c, d** Simulated community dynamics with 900 species where each strain has a unique combination of allocation parameters *f*_r_ and 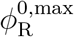. A single strain ‘wins’, with intermediate allocation values in both the P and R sectors. **e, f** Pairwise invasibility plots constructed for both allocation parameters predict a single optimal strategy that is globally uninvasible. **g** Simulated evolutionary trajectories with random mutations match the trajectory predicted by adaptive dynamics.

We simulated eco-evolutionary dynamics in two ways. First, we started with a large pool of many competing strains, each with randomly generated distinct allocation strategies (Methods). By simulating these dynamics we observed that a single surviving strain eventually emerged following a period of intense competition (Fig. 2c,d). The winning strategy had intermediate values of *f*_r_ = 0.55 and 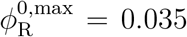. Second, we simulated evolutionary dynamics starting from a single strain with no pre-allocation, i.e., *f*_r_ = 1 and 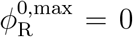. We let this strain grow, and then successively introduced new randomly mutants of the current resident strain one at a time (Methods). New mutants could outcompete their parents and take over the community, becoming parents for the next mutant. Interestingly, we found that after many successive mutations, the simulations repeatably converged to the same winning strategy (Fig. 2g, red), suggesting that it is evolutionarily accessible, stable, and robust to the use of different simulation methods. This winning strategy resembles that of a generalist and best balances growth rates with lag times across both rich and poor environments. On the other hand, strategies that over-invested to become specialists in either ribosomal or metabolic enzyme reserves were outcompeted because: while they did better in the presence of one nutrient (rich and poor, respectively), they fared much worse in the presence of the other.

To understand this outcome theoretically, we used the mathematical framework of adaptive dynamics [42, 43, 37] (see Methods for details). This framework successfully predicted not only the winning strategy, but also the evolutionary trajectory that led to it (Fig. 2g, blue). To gain further insight, we performed a pairwise invasibility analysis, where we measured the invasion growth rate of each allocation strategy in the presence of every other strategy, when present as the lone resident (Fig. 2e,f shows plots for both allocation parameters *f*_r_ and 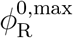). This analysis showed that the winning strategy could successfully invade every other allocation strategy, and could not be invaded by any other strategy, providing theoretical confirmation that the winner was indeed the evolutionarily stable strategy (ESS) for this pattern of environmental fluctuations. Together, these results show that symmetric environments selected a single ESS corresponding to a unique growth-rate profile.

### Emergence of two coexisting strains in asymmetrically fluctuating environments

Next, we ask how changing the balance of nutrient supply alters evolutionary outcomes. To do this, we repeated the same serial-dilution simulations but imposed an asymmetric environment in which the rich nutrient was supplied in excess, *c*_r_*/c*_p_ = 100 (Fig. 3a,b).

**Figure 3.**
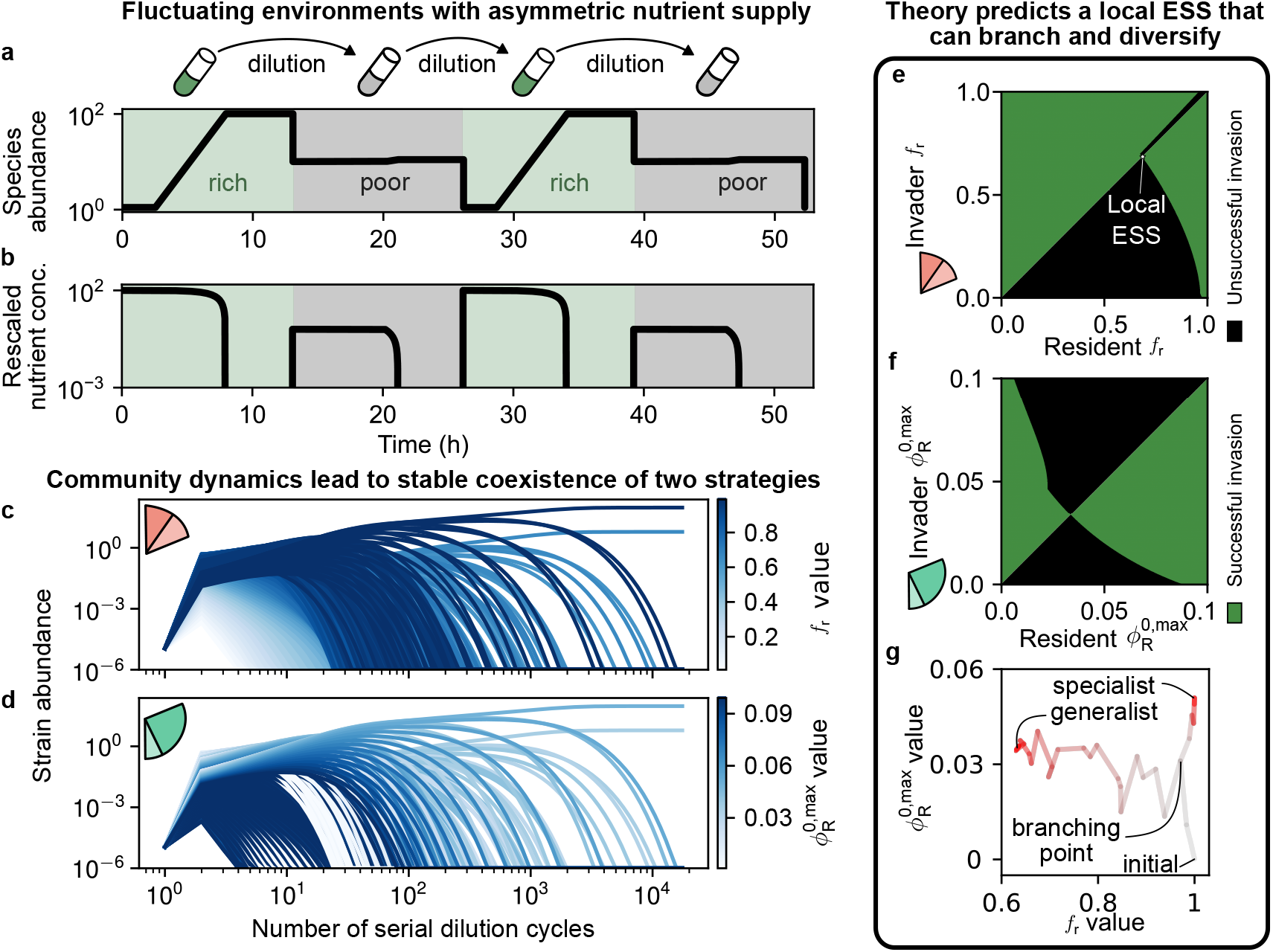
Asymmetric environments allow for the stable coexistence of two strains. **a, b** Example dynamics as in Figure 2a and b. Here, nutrient concentration ratio, *c*_r_*/c*_p_ = 100 and dilution factor *D* = 10. **c, d** Simulated community dynamics as in Figure 2. A specialist on the rich nutrient, with high *f*_r_ and 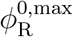 coexists with a generalist with lower allocation values in both the P and R sectors. **e, f** Pairwise invasibility plots show that a locally stable strategy in *f*_r_ can invade and be invaded by mutant strategies, suggesting the possibility of branching. **g** An example evolutionary trajectory shows the branching process where one strain eventually becomes the specialist (with high *f*_r_) and the other the generalist.

In contrast to symmetric environments, community dynamics no longer converged to a single surviving strain. Instead, starting from a large pool with many randomly generated strains, competition led to the stable coexistence of two strains with distinct allocation strategies (Fig. 3c,d). One strain adopted high values of both *f*_r_ = 0.99 and 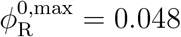, corresponding to a specialist on the rich nutrient that prioritized current growth with a significant fraction of inactive ribosomes. The other strain maintained lower values of both parameters *f*_r_ = 0.65 and 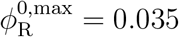, corresponding to a generalist that sacrificed growth rates on the rich nutrient to reduce lag times and perform better on the poor nutrient.

Thus surprisingly, these two strains could stably coexist even in these extremely asymmetric fluctuating environments where the poor nutrient was supplied at much lower levels. Naively, as nutrient supply shifts from symmetric to asymmetric environments, we might expect the winning strategy to smoothly shift as well. Instead, our results show that as environments become more asymmetric, selection may favor diversification. No single allocation strategy can simultaneously adapt to maintain high growth rates in the rich nutrient as well as low lag times during transitions to poor conditions. This is because of the growth rate–lag tradeoff (Methods). Thus, a strategy that grows fast on the rich nutrient faces a long lag time on the poor nutrient, which leaves open the poor nutrient niche. A new specialist strain can thus evolve to take advantage of this open niche. This explains why such stable coexistence can reproducibly and spontaneously arise in asymmetric environments.

We could also recapitulate this diversification theoretically using adaptive dynamics (Methods and SI). Pairwise invasibility analysis showed that the branching arose due to the metabolic enzyme pre-allocation *f*_r_. We observed a point 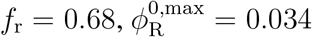 that was a local ESS (Fig. 3e). Although it was immune to invasion by strains differing by an infinitesimal amount (Fig. 3e, black close to star); larger mutations could introduce strains that would invade this strain and coexist with it (Fig. 3e, green region above star). Once they coexisted, selection allowed them to diverge in their metabolic enzyme pre-allocation *f*_r_, which led to the strains diversifying while continuing to coexist. These changes in *f*_r_ ultimately also affected each strain’s selection on inactive ribosomes 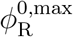 (Fig. 3f).

Stochastic evolutionary simulations confirmed this prediction. Starting from a single ancestor, the population eventually split into two lineages that diverged toward the specialist and generalist strategies we observed in our previous simulations (Fig. 3g and Fig. 3c,d). Importantly, these coexisting strains emerged clonally from a single lineage through evolutionary dynamics in the asymmetric environment. Together, these results show that asymmetric nutrient supply can reproducibly promote the coexistence of strains with distinct growth-rate profiles even in rapidly fluctuating environments.

### Mapping environmental fluctuations to evolved allocation strategies

We showed that evolutionary outcomes are strongly dependent on environmental conditions. To systematically establish this dependence, we varied the dilution factor *D* and the nutrient supply ratio *c*_r_*/c*_p_, and mapped the resulting evolved strategies across environments (Fig. 4).

**Figure 4.**
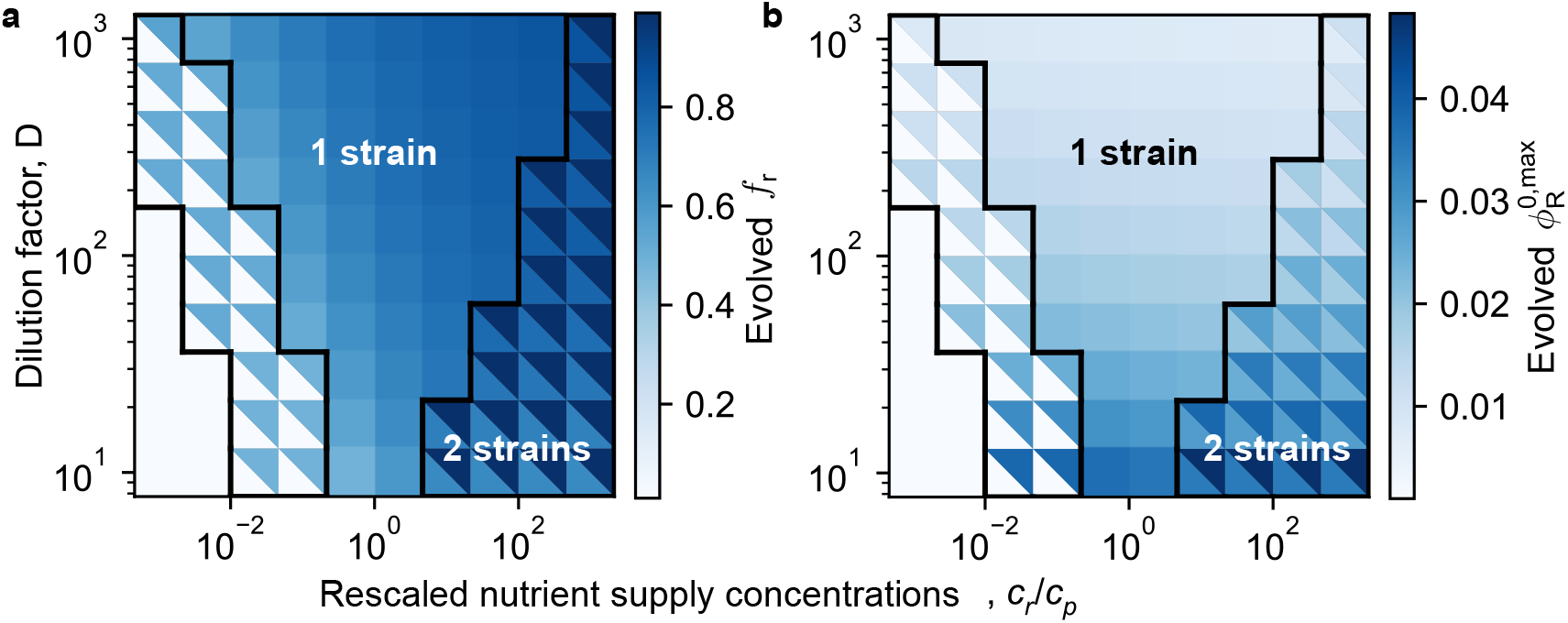
Evolutionarily stable strategies change predictably with different environmental fluctuation patterns. Heatmaps of the allocation strategies of evolved strains in community simulations as a function of environmental parameters: dilution factor *D* and relative supply concentration of rich to poor nutrients *c*_r_*/c*_p_; shown are the two evolved allocation parameters *f*_r_ (in **a**) and 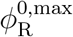 (in **b**). Black borders separate regions where only a single strain is evolutionarily stable from regions where a pair of strains coexists.

We found that environments separated into two qualitatively distinct outcomes: those where evolution consistently led to a single dominant strain, and others where it resulted in the stable coexistence of two strains with different allocation strategies (Fig. 4a–b). Specifically, symmetric environments where rich and poor nutrients were equally likely to be available selected for a single strain, while asymmetric environments that were skewed towards the availability of either the rich or poor nutrient selected for two coexisting strains. Moreover, a larger dilution factor *D* broadened the region supporting a single winning strain.

The key factor shaping which strategy was evolutionarily stable in each environment was the growth–lag tradeoff. In each environment, growth and lag were selected for in different ways. In environments with larger dilution, populations spend a longer time growing until resource depletion (see SI), making the growth phase more relevant than the lag one. Thus, a larger *D* selected for more growth. Asymmetric environments on the other hand selected against long lag times during both nutrient upshifts and downshifts. Therefore, as environments became more asymmetric, strains could not simultaneously have short lags in both nutrient shifts, which promoted divergence into two strains. In environments skewed towards greater availability of the rich nutrient (*c*_r_*/c*_p_ ≫1), we observed the emergence of a near-specialist strain that derived most of its growth from the rich nutrient, and a near-generalist strain that grew relatively poorly on the rich nutrient but derived a larger fraction of its growth from the poor nutrient. In contrast, in environments skewed towards greater availability of the poor nutrient (*c*_r_*/c*_p_ ≪1), the two coexisting strains instead comprised a near-specialist favoring the poor nutrient (*f*_r_≈0) and a near-generalist deriving a larger fraction of its growth from the rich nutrient (*f*_r_ ≈0.5).

A key and counterintuitive result was that these strategies did not simply follow nutrient quality. For example, once environments became sufficiently skewed so that the rich nutrient was hardly available (*c*_r_*/c*_p_ *<* 10^*−*2^ in Fig. 3a), this latter near-generalist could no longer support enough of its growth on the rich nutrient, and we observed another transition to a single dominant strategy, that of the near-specialist strain on the poor nutrient. Notably, this evolved strain would be a slow-grower on the rich nutrient. Yet, this slow-growing strain outcompetes faster-growing mutants arising in these environments (Fig. 3, white region). This occurs because fast-growing strains pay a cost for maintaining ribosomes and metabolic capacity that outweighs the benefit from growth on a meager concentration of the rich nutrient. Hence, the pattern of nutrient availability in the environment can counteract nutrient quality during selection of growth-rate profiles.

Taken together, these results demonstrate that distinct patterns of environmental fluctuations can rapidly select for different growth-rate profiles among closely related strains, even with the same underlying enzymatic repertoire, purely through how strains pre-allocate their proteome towards different nutrients.

### Evolved strains spanning environmental histories recapitulate patterns in data from natural isolates

The results above demonstrate that environmental fluctuations between two nutrients can generate growth-rate variability and strain coexistence. However, natural environments are more complex: bacteria encounter multiple carbon sources whose availability changes unpredictably over time. We therefore asked whether our results extend to settings with more nutrients and less predictable fluctuations.

To address this, we generalized our model to four nutrient media defined by two carbon sources and the presence or absence of amino acids (Fig. 5a). Two media were rich (each sugar supplemented with amino acids) and two were poor (each sugar alone as the unique carbon source), to capture several different kinds of nutrients that might become available in natural environments. We also relaxed our previous assumption that media always alternate from rich to poor and back by allowing for stochastic transitions: at each dilution step, the next medium was drawn randomly from other media (Methods). Importantly, transitions between some states were more likely than others, and thus each environment could be represented as a state transition network with its own pattern of environmental fluctuations (Fig. 5a). In any environment, strains that could anticipate which nutrient will appear next by appropriately adjusting their allocation strategy would naturally have a higher long-term growth rate and outcompete others.

**Figure 5.**
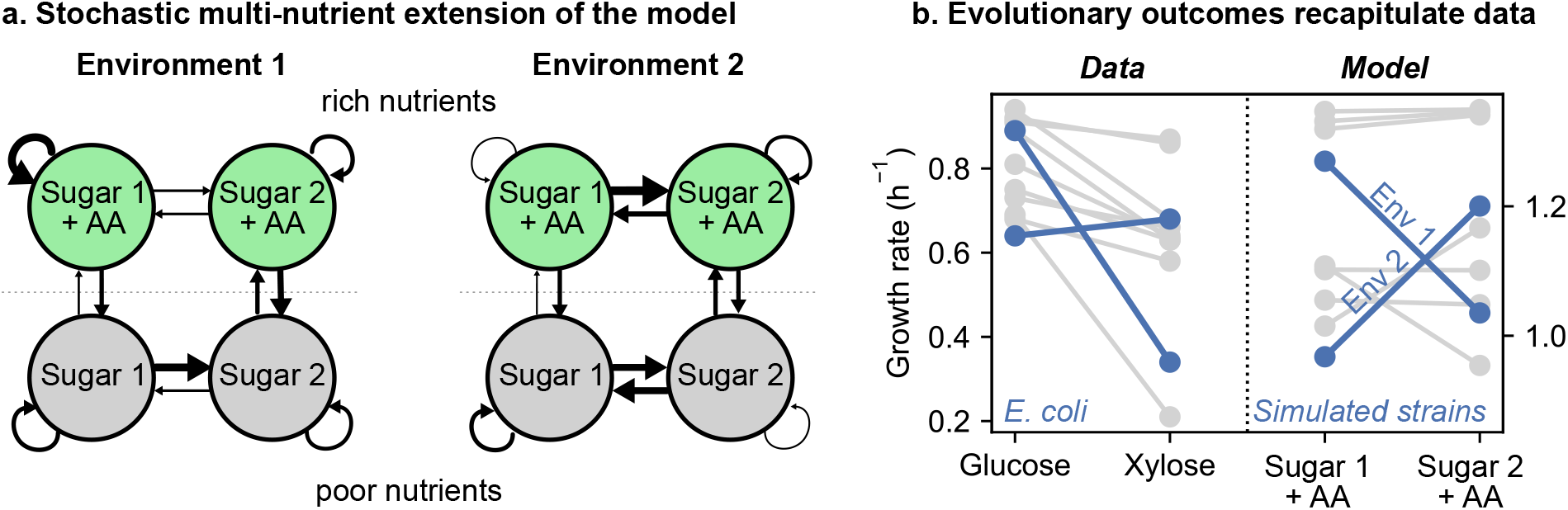
Evolved strains spanning environmental histories recapitulate patterns in data from natural isolates. **a** We extended our model to include 4 distinct nutrient media, 2 being rich and 2 being poor. Each medium had one of two randomly generated sugars (Methods); rich media also had amino acids (AA) in addition. Between two serial dilution cycles, we changed the medium randomly according to the transition diagram shown. To simulate a variety of environmental fluctuation patterns, we varied the transition probabilities between different nutrients (see Methods for details). We evolved strains in each environment and surveyed isolates, much like during laboratory isolation. **(b)** Growth rates of evolved strains in different media from our model recapitulate variability and patterns observed in experimental data; for comparison we also show *E. coli* strains from Fig. 1a.

We then simulated evolution across a range of environments with different transition patterns (Fig. 5b). In some environments, it was more likely to switch from sugar 1 to 2 but not back (Fig. 5a, environment 1); in others it was instead very likely to switch from sugar 2 to 1 (Fig. 5a, environment 2). In each environment, we evolved strains independently using the stochastic mutation–selection dynamics described above. We then collected all the resulting isolates evolved across environments with distinct transition patterns, analogous to surveying natural strains sampled from distinct habitats.

The growth-rate profiles of these evolved strains, measured on the two rich media, reproduced the key qualitative features observed in conspecific isolates of natural bacteria (Fig. 5c). Strains from different environmental histories exhibited a several-fold variation in growth rate on the same medium, consistent with the *E. coli* data in Fig. 1a. Crucially, growth rate crossovers emerged spontaneously: strains exposed to environments that often transitioned to sugar 1 after sugar 2 became available evolved to be slow-growers in media containing sugar 2. Similarly, strains exposed to the opposite environmental transition patterns rapidly evolved to become slow-growers on sugar 1 instead. These inversions did not require any differences in metabolic enzymes between strains: they arose solely from differences in proteome allocation shaped by each strain’s environmental history.

This pattern provides a possible mechanistic resolution to the puzzle regarding strain growth rate data motivating this work. Growth rate differences across resource do not necessarily reflect biochemical properties, but rather statistical properties of environmental variability. A strain that grows faster on a conventionally poorer nutrient might have evolved in an environment where that nutrient was supplied more abundantly, selecting for an allocation strategy poised to exploit it. In this view, the growth-rate inversions observed across conspecific isolates (Fig. 1a) are a natural consequence of evolution in environments with distinct fluctuation patterns, and the growth-rate profile of a strain serves as a record of the environments in which it has evolved.

## Discussion

Why do closely related bacterial strains often grow at very different rates on the same nutrient? Standard explanations point to nutrient quality and to differences in metabolic enzymes [5, 3]. Our results show that these explanations rooted in biochemical properties and constraints are not necessary to explain the observed variability. Even when strains share the same metabolic repertoire, selection in fluctuating environments can generate distinct growth-rate profiles by favoring different patterns of proteome allocation [23, 31, 24].

Our central idea is that cells must adjust their proteome allocation between growing now and preparing for what comes next. A strain can maintain inactive ribosomes that help it respond rapidly to nutrient upshifts, but this comes at the cost of reduced growth in rich medium [31, 32]. Likewise, a strain can invest strongly in enzymes for rich nutrients and grow rapidly when those nutrients are present, but then faces a longer delay after a transition to poorer conditions [23]. These tradeoffs mean that no single allocation strategy is best in every phase of a fluctuating environment [24, 25]. Selection therefore acts on cumulative growth over an entire sequence of nutrient shifts, rather than on growth rate in any one medium [33, 34].

Our result provides an explanation of why the evolutionarily selected strain need not be the fastest grower in rich medium. In environments with low effective mortality and a relatively larger contribution from poor nutrients, evolution selects allocation strategies that grow comparatively slowly on the rich nutrient. This does not mean that slow growth is intrinsically beneficial. Instead, it reflects the fact that these strategies pay lower costs over the full fluctuation cycle and perform better when poor conditions recur. A growth measurement in a single medium can therefore give a misleading picture of fitness: the fastest strain at one instant need not achieve the largest long-term growth [23, 24].

Our results also show that changing the fluctuation pattern can change the qualitative outcome of evolution. In symmetric environments, competition selects a single allocation strategy that balances performance across rich and poor conditions. In contrast, across a region of asymmetric environments, this compromise becomes unstable. One lineage evolves toward rapid growth in the rich phase, while another retains greater preparedness for the poor phase. These two strains coexist because each performs well during a different part of the cycle. Thus, temporal variation can generate strain-level ecological differentiation even without spatial structure, cross-feeding, or differences in the metabolic enzyme repertoire [35, 36].

The multi-nutrient model shows that this mechanism can also generate growth-rate patterns resembling those observed across natural isolates [3]. In particular, strains evolving under different nutrient-transition statistics develop growth-rate crossovers: a strain can grow slowly in one rich medium and rapidly in another, despite having the same underlying metabolic capabilities as competing strains. This suggests that growth-rate profiles may contain information about the fluctuation regimes under which a lineage evolved [21, 22]. Such information will not uniquely identify environmental history, since distinct environments may select similar phenotypes. Nevertheless, growth measurements across multiple nutrients could help constrain the ecological histories that remain compatible with a strain’s phenotype.

These results make several experimentally testable predictions. Evolutionary experiments that independently vary dilution factor and the relative supply of rich and poor nutrients should select distinct allocation strategies, together with distinct combinations of growth rates and lag times. The model predicts regions in which the selected strain grows relatively slowly in rich medium, as well as regions in which initially clonal populations diversify into two stable strategies. Measuring growth rates alone will not be sufficient to test this mechanism. The decisive signature is the associated pattern of lag times across nutrient upshifts and downshifts [23, 31]. While our model was deliberately simplified in order to highlight our central idea, several extensions of it are important in order to study more realistic ecological settings. We have represented the proteome using a coarse-grained sectors and treated allocation parameters as heritable traits under selection. Cells implement this proteome allocation through regulatory networks, protein turnover, and metabolic feedbacks [29, 6], and including some of these details may introduce additional tradeoffs or quantitatively affect the growth rate variability we observed. Further, including cross-feeding and more complex environments may introduce multiple nutrients in every growth cycle, creating further routes to coexistence and help us study these dynamics in more diverse ecological communities [35, 44]. Our results establish nevertheless a general point: fluctuating environments can generate substantial growth-rate variation and stable strain-level diversity through selection on proteome allocation alone.

More broadly, our results suggest that growth rate is not simply a property of an organism– nutrient pair. It also depends on the history of environmental fluctuations that shaped the strain’s allocation strategy. A slow grower on a specific nutrient might have evolved in a history of environments with lower availability of this nutrient [6]. These observations might ultimately help explain the substantial variability in growth rates in strains of the same species that share identical core metabolic and biosynthetic genes [11, 3], especially such as *E. coli* in the human gut [45]. Here too, different individuals may carry different strains of the same species [46], shaped in part by the differences in their gut environments and dietary habits. Our work suggests the possibility that many of these strain-level differences may be explained by rapid evolution of proteome allocation in these different patterns of environmental fluctuations.

## Acknowledgements

We are grateful to Jonas Cremer for valuable discussions and comments on the manuscript. This research was supported in part by the International Centre for Theoretical Sciences (ICTS) for participating in the program - Unifying Theories in High-Dimensional Biophysics (code: ICTS/UHDB2025/07). A.G. acknowledges support from the Ashok and Gita Vaish Junior Researcher Award, the Govt. of India’s Ramanujan Fellowship, the Centre for Artificial Learning and Intelligence for Biological Research and Education, ICTS-TIFR, as well as the DAE, Govt. of India, under project no. RTI4001. A.M.D acknowledges support from the ICTS Long Term Visiting Students Program, 2025. A.G. and A.M.D also acknowledge support from the ICTP through the Associates Programme and from the Simons Foundation, whose grant (Record ID: SFI-MPS-T-Institutes-00012057, AD) also supported this work. J.G. acknowledges the support of Fondo Italiano per la Scienza - FIS (CUP J53C23002290001).

## Data and code availability

All data and code used in this paper is available as a GitHub repository at the following link: https://github.com/arjun-projcode/Code-envhist

## Methods

### Modelling bacterial growth

We work in the framework of [5], where the proteome is divided into three sectors quantifies by the fraction of protein mass devoted to a coarse-grained function. The three sectors we consider are: the metabolic sector *ϕ*_P_, the ribosomal sector *ϕ*_R_, and the housekeeping sector *ϕ*_Q_, with *ϕ*_P_ + *ϕ*_Q_ + *ϕ*_R_ = 1. We consider two distinct phases of growth: steady state (exponential) growth phase and a lag phase. During steady state exponential growth, the growth rate equals both the nutrient influx and outflux determined by the P and R sectors respectively [5, 6, 32],

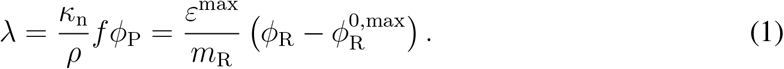

The first term is the flux from metabolic enzymes, with *κ*_n_ denoting the nutrient quality, *f* denoting the fraction of enzymes allocated to current growth, and *ρ* being a conversion factor from RNA/protein ratio to proteome fraction. The second term represents the flux due to ribosomes, where *ε*^max^ is the maximal elongation rate of each ribosome; *m*_R_ the mass of each ribosome, and 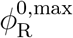 the maximal fraction of inactive ribosomes (see [32] and SI).

We initially consider two types of nutrients: rich and poor. The lag times are the effective time lost due to sector re-allocation between different nutrients, thus 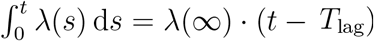, giving

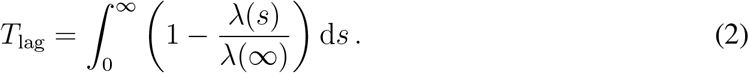

Note that *λ*(*s*) is the instantaneous growth rate —*λ*(*t*) = *d* log *B*(*t*)*/dt* where *B*(*t*) is the total biomass at time t — determined by the proteome allocation at time *s*. The entire time period of reallocation contributes to the lag time. Upshifts require synthesis of the R sector whereas downshifts require P sector synthesis. Note that we imagine a situation where the additional metabolic enzymes required after nutrient upshift are not limiting to growth, and therefore set *f*_p_ = 1. We assume logistic growth of the limiting sector: *ϕ*?(*t*) = *ϕ*(*t*)(*ϕ*_final_ *− ϕ*(*t*)). Differentiating Equation 1, substituting these dynamics and plugging into Equation 2 gives us the lag time,

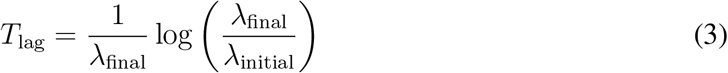

where *λ*_final_ is the steady-state post-shift growth rate and *λ*_initial_ is the effective initial growth rate due to pre-allocation. For nutrient upshifts, we assume that ribosomes immediately elongate at the rich nutrient steady-state rate and that the fraction of inactive ribosomes is that at rich nutrient steady-state. This assumption holds since the amount of the alarmone ppGpp, which regulates ribosome activity, can change on a timescale of 5-10 minutes [32]. Substituting from we [8] derive

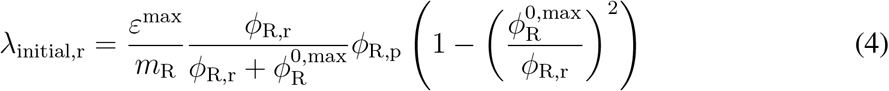

Where *ϕ*_R,r_ and *ϕ*_R,p_ are the steady-state R sector allocations in rich and poor conditions respectively. For downshifts, we assume the metabolic enzymes that are pre-allocated determine initial growth. Thus

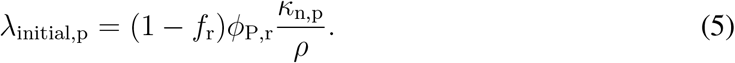

### Well-adapted strategies in fluctuating environments

We assume a model of batch dynamics where growth occurs in a nutrient condition for a fixed time *T*, chosen to be sufficiently large such that nutrient depletion occurs, following which the culture is diluted and transferred to another nutrient. Fluctuations in the environment occur by alternating between the rich and poor nutrients. Two environmental parameters define an environment: the dilution factor, *D* and the relative concentration of the rich nutrient (*c*_r_*/c*_p_). For a single strain growing in this environment with biomass *M* (*t*), dynamics are given by 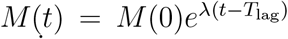. Assuming nutrient depletion occurs proportional to growth as 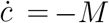, we define the depletion time of each nutrient as t. Then, with a resident strain at steady-state in a fluctuating environment (i.e., the growth across both nutrients matches dilution), an invader strain successfully invades if its growth across the rich and poor nutrients is higher than dilution. This is done in the environment of the resident, i.e., the depletion time of either nutrient is solely determined by the parent. Then we can derive the invasion fitness as

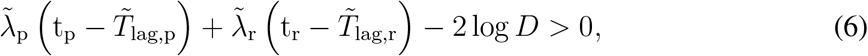

where terms 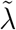and 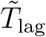 are growth rates and lag times of the invader respectively.

### Details of simulations

In our model, we fix all parameters except 2 allocation parameters, *f*_r_ and 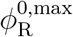 and 2 environmental parameters, *D* and *c*_r_*/c*_p_. The maximal elongation rate, *ε*^max^ is a fixed physiological property for a species, and the size of the housekeeping sector, *ϕ*_Q_ is growth independent. Further, we do not aim to examine the effect of varying nutrient qualities for either rich or poor nutrients in this study. For a full list of parameters, see SI. In order to simulate community dynamics with multiple strains, we created a grid of strains with *f*_r_ ∈ (0.01, 0.99) and 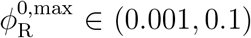. This gives each strain a unique growth rate and lag time on either nutrient. All strains were initialized at an abundance of 10^*−*5^. We defined the growth function

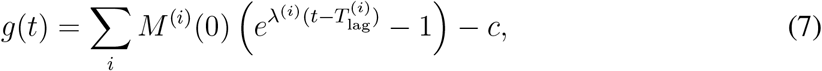

whose roots give us the depletion time of the community, t_dep_, where *i* represents each strain.

Then the total growth of strain *i* is given by 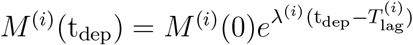.

To construct pairwise invasibility matrices, we computed the roots of the gradient of the invasion fitness defined by Equation 6 in the two allocation parameters for a given set of environmental parameters. The roots give us evolutionarily singular points [42], and we then constructed invasibility matrices by computing Equation 6 for every resident-invader pair in either allocation parameter, keeping the other fixed at the singular value.

We initialized a resident population with *f*_r_ = 1 *−* 10^*−*4^ and 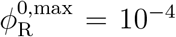. Mutants had 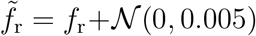and 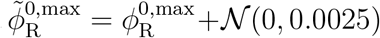 We introduced mutants sequentially, waiting until the ecological dynamics reached steady state before introducing another mutant. To obtain the optimal trajectory predicted by adaptive dynamics, we numerically computed the direction of highest invasion fitness in 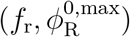 space. This corresponds to the canonical equation of adaptive dynamics,

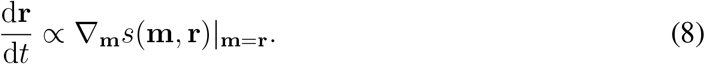

Note that *s*(**m, r**) is defined by Equation 6.

In order to generate the heatmaps in Figure 4, we simulated the community dynamics described above in a range of environments defined by varying *D* and *c*_r_*/c*_p_ ratios. The remaining strains were then invaded by the numerically computed ‘optimal’ strains, given by the roots of the gradient of Equation 6, and dynamics were allowed to play out. The ‘winners’ of this exercise were then used to populate the cells in Figure 4.

### Extension of the model

We consider an extension of the framework to a multi-nutrient setup where there are two rich nutrients and two poor nutrients. We assume that the nutrient conditions are chosen such that achieving steady-state growth after an upshift is limited by the synthesis of ribosomes only. For instance, this is the case when each poor and rich pair correspond to the same carbon source in either the presence or absence of supplemented amino acids. To maintain a distinction between *ϕ*_R_-limited transitions and *ϕ*_P_-limited transitions, we assume that transitions between nutrients are restricted to only three types: the same nutrient repeats (self-transition), the environment switches to a different carbon source of the same quality (limited by synthesis of *new* metabolic enzymes in *ϕ*_P_), or the environment switches to rich/poor with the same carbon source (upshifts are limited by *ϕ*_R_, downshifts are limited by synthesis of *more of the same* metabolic enzymes within *ϕ*_P_). See Figure 5a for an overview.

Note that this is the minimal extension to the current setup; and can be expanded to include multiple carbon sources of varying nutrient qualities. We assume all nutrients arrive in equal quantities; *c*_nutrient_ = 1. We randomly sample transitions between nutrients as p_transition_ ∝ 10^*k*^ where *k* ∼*U* (−3, 3), with the appropriate normalization. We set *κ*_n_*/ρ* for both rich nutrients to 5.3, and that of poor nutrients to 0.2. We allowed different strains to grow and evolve in such randomly sampled environments for 1000 mutation steps, and the evolved strains were surveyed with their final growth rates plotted in Figure 5b.

## Notes

https://github.com/arjun-projcode/Code-envhist

### Competing Interest Statement

The authors have declared no competing interest.

