## Supplementary Information for "Environmental history can counteract nutrient quality in shaping the evolution of bacterial growth rates"

This file contains:

Supplementary Figures 1-3
Supplementary Text

### Supplementary Figures

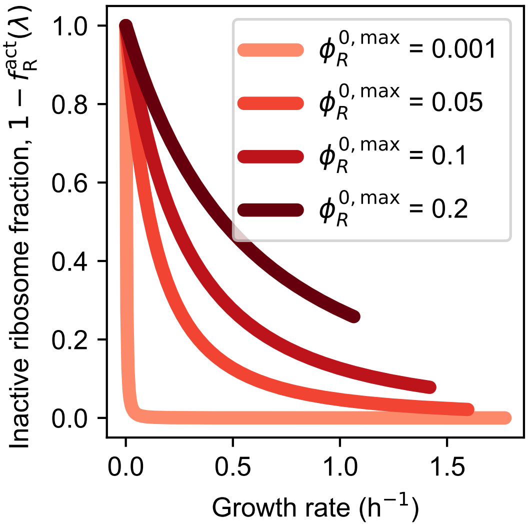


Supplementary Figure 1: The fraction of inactive ribosomes as a function of growth rate (given by Equation S.18), plotted for various values of the parameter $\phi_{\mathrm{R}}^{0,\max}$. Observe that though $\phi_{\mathrm{R}}^{0,\max}$ is constant, $f_{\mathrm{R}}^{\mathrm{act}}$ is a growth-rate dependent term.

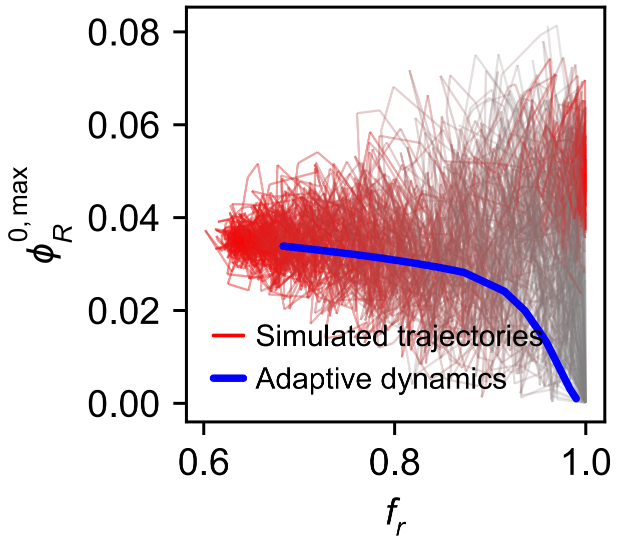


Supplementary Figure 2: 128 trajectories of branching evolution plotted for the environmental parameters $D = 10$, $c_r/c_p = 100$ (see Figure 3g). Shown in red are various evolutionary trajectories in trait space, starting from initial values of $f_r \sim 1$ and $\phi_R^{0,max} \sim 0$. Trajectories get darker as time progresses, and display a branching into two strains. Notably, the branching happens at a different point in trait space for each trajectory. Shown in blue is the evolutionary trajectory predicted by adaptive dynamics (see Methods).

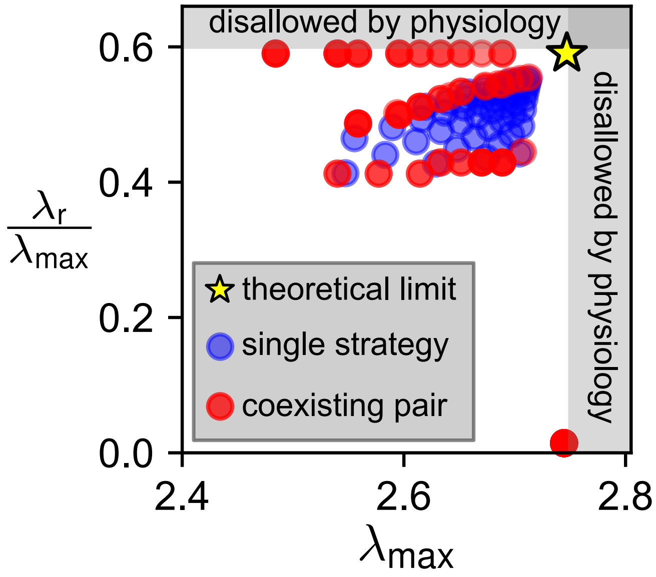


Supplementary Figure 3: We take evolved strategies (in $\phi_R^{0,max}$ and $f_r$) from Figure 4 and compute $\lambda_r$ and $\lambda_{max}$ for each of them (see Equation S.7 and Equation S.9). This creates a 'physiology space' that allows us to plot the distinct strategies that evolve in different environments. The strains with very high $\lambda_r/\lambda_{max}$ are rich nutrient specialists; while those with higher $\lambda_{max}$ (and low $\lambda_r/\lambda_{max}$) are poor nutrient specialists. No environment selects for the theoretically 'best' strain (star). Blue points represent evolved strains which survive as the only strain in their environment, while red points represent cases where evolution leads to coexisting strain pairs.

### Supplementary Text

#### Model

We follow the growth laws of Ref. [5]. We assume the population to be homogeneous within each strain with no spatial or age structure, and that growth happens in 2 distinct phases: steady-state growth, and lag. We initially consider 2 nutrients, rich and poor (for nutrient qualities, see Table S1 at the end of this section). At steady-state growth,

$$\lambda = \frac{\kappa_{\mathrm{n}}}{\rho} f \phi_{\mathrm{P}}, \tag{S.1}$$

where $\lambda$ is the growth rate, $\kappa_{\mathrm{n}}$ the nutrient quality and $f$ is the fraction of the P sector allocated to current growth. In this model we will assume that $f$ for the poor nutrient is 1, i.e., cells allocate 100% of their P-sector resources to growth in the poor nutrient. This is because we consider nutrient upshifts (i.e., shifts from the poor to rich nutrient) to be limited by inadequate R sector flux, thus there is no benefit to pre-allocation in the P sector. Note that in a further extension to this model (see Figure 5 and Methods), we consider multiple carbon sources: in that case, pre-allocation exists in the P sector during growth on poor nutrient conditions since transitions between two poor nutrients of the same quality would require different metabolic enzymes.

Ref. [6] shows that the elongation rate per ribosome, $\varepsilon$ and inactive ribosome fraction, $\phi_{\mathrm{R}}^{0}$ are functions of the growth rate. The cell balances an inactive reserve ribosome fraction with the need to maintain per-ribosome elongation rates. They derive

$$\lambda = \frac{\varepsilon(\lambda)}{m_{\mathrm{R}}} \left(\phi_{\mathrm{R}} - \phi_{\mathrm{R}}^{0}(\lambda)\right) = \frac{\varepsilon^{\max}}{m_{\mathrm{R}}} \left(\phi_{\mathrm{R}} - \phi_{\mathrm{R}}^{0,\max}\right), \tag{S.2}$$

where $m_{\mathrm{R}}$ is the mass of a ribosome, and $\phi_{\mathrm{R}}^{0,\max}$ is the maximal fraction of inactive ribosomes, i.e., $\phi_{\mathrm{R}}^{0}(\lambda = 0)$. We denote by $\phi^{\max}$ the maximum possible size of the R sector (i.e., the limiting case where no P sector is needed to produce a nutrient influx),

$$\phi^{\max} = \phi_{\mathrm{R}} + \phi_{\mathrm{P}} = 1 - \phi_{\mathrm{Q}}. \tag{S.3}$$

By substituting $\phi_{\mathrm{P}}$ from Equation S.1 and $\lambda$ from Equation S.2, one can derive the total R sector allocation in a given medium,

$$\phi_{\mathrm{R}} = \frac{\varepsilon^{\max}\rho\phi_{\mathrm{R}}^{0,\max} + \kappa_{\mathrm{n}} m_{\mathrm{R}} f \phi^{\max}}{\varepsilon^{\max}\rho + \kappa_{\mathrm{n}} m_{\mathrm{R}} f}. \tag{S.4}$$

Using Equation S.3,

$$\phi_{\mathrm{P}} = \left(\phi^{\max} - \phi_{\mathrm{R}}^{0,\max}\right)\left(\frac{\varepsilon^{\max}\rho}{\varepsilon^{\max}\rho + \kappa_{\mathrm{n}} m_{\mathrm{R}} f}\right) \tag{S.5}$$

gives us the total $P$ sector allocation in a particular medium. Finally we can use Equation S.1, Equation S.2 and Equation S.3 to get

$$\lambda = \left(\phi^{\max} - \phi_{\mathrm{R}}^{0,\max}\right)\left(\frac{m_{\mathrm{R}}}{\varepsilon^{\max}} + \frac{\rho}{\kappa_{\mathrm{n}} f}\right)^{-1}. \tag{S.6}$$

Equation S.6 essentially defines the growth rate as a function of various physiological parameters. We can define the maximum possible growth rate of the cell,

$$\lambda_{\max} = \left(\phi^{\max} - \phi_{\mathrm{R}}^{0,\max}\right)\frac{\varepsilon^{\max}}{m_{\mathrm{R}}}. \tag{S.7}$$

This represents the limit where all ribosomes are active, and there is sufficient nutrient influx for the cell to allocate all possible proteome resources to ribosomes. However, this 'true optimum' is hard to achieve, and we can define the maximal possible growth rate in a particular medium. This is achieved when the P sector allocation $f = 1$:

$$\lambda^* = \left(\phi^{\max} - \phi_{\mathrm{R}}^{0,\max}\right)\left(\frac{m_{\mathrm{R}}}{\varepsilon^{\max}} + \frac{\rho}{\kappa_{\mathrm{n}}}\right)^{-1}. \tag{S.8}$$

Equation S.7 sets a theoretical upper bound on the growth rate that is determined by $\phi_{\mathrm{R}}^{0,\max}$ and not $f_{\mathrm{r}}$. We can calculate the ratio $\lambda_{\mathrm{r}}/\lambda_{\max}$ using Equation S.6 and Equation S.7,

$$\frac{\lambda_{\mathrm{r}}}{\lambda_{\max}} = \frac{m_{\mathrm{R}}}{\varepsilon^{\max}}\left(\frac{m_{\mathrm{R}}}{\varepsilon^{\max}} + \frac{\rho}{\kappa_{\mathrm{n}} f}\right)^{-1}. \tag{S.9}$$

This ratio depends only on $f_{\mathrm{r}}$, and therefore we can create a phase space using $\lambda_{\max}$ and $\lambda_{\mathrm{r}}/\lambda_{\max}$. This is analogous to the $f_{\mathrm{r}}$ and $\phi_{\mathrm{R}}^{0,\max}$ phase space, however it only includes physiological growth rate properties that can be experimentally measured.

Given a nutrient condition, we can determine physiological bounds for these quantities. First, $\lambda_{\max}$ is maximized when $\phi_{\mathrm{R}}^{0,\max} = 0$. For the parameter values in Table 1, we find that this occurs at $\lambda_{\max} = 2.75\ \mathrm{h}^{-1}$. Similarly $\lambda_{\mathrm{r}}/\lambda_{\max}$ is maximized when $f_{\mathrm{r}} = 1$, or $\lambda_{\mathrm{r}}/\lambda_{\max} = 0.6$. In Supplementary Figure 3 we plot the various optimal strains from Figure 4 in this space, with physiological limits thus calculated marked as bounds.

**Nutrient transition kinetics**

Although nutrient transitions involve the expression of many different proteins ([3, 6]), we assume that lag times depend only on the limiting sector. Thus upshifts are limited by $\phi_{\mathrm{R}}$ and downshifts by $\phi_{\mathrm{P}}$. This arises from differing allocation dictated by Equation S.4 and Equation S.5. We calculate lag time as the effective time lost due to reallocation dynamics,

$$\int_0^t \lambda(s)\,\mathrm{d}s = \lambda(\infty)\cdot(t - T_{\mathrm{lag}}), \tag{S.10}$$

where $t = \infty$ is at steady-state growth. Here the right hand side represents the effective 2 phase model of growth. Rearranging, we get

$$T_{\mathrm{lag}} = \int_0^\infty \left(1 - \frac{\lambda(s)}{\lambda(\infty)}\right)\mathrm{d}s\,. \tag{S.11}$$

During upshifts kinetics are determined by R sector production. Let $\phi_{\mathrm{R,r}}$ be the steady-state R sector in the rich medium. We assume logistic growth for the R sector:

$$\begin{aligned}\frac{\mathrm{d}\phi_{\mathrm{R}}(t)}{\mathrm{d}t} &= \frac{\varepsilon(t)}{m_{\mathrm{R}}}\phi_{\mathrm{R}}^{\mathrm{act}}(t)\left(\phi_{\mathrm{R,r}} - \phi_{\mathrm{R}}(t)\right) \\ &= \lambda(t)\left(\phi_{\mathrm{R,r}} - \phi_{\mathrm{R}}(t)\right),\end{aligned} \tag{S.12}$$

where the second equality follows from Equation S.2. We can differentiate Equation S.2 to get

$$\frac{\mathrm{d}\lambda(t)}{\mathrm{d}t} = \frac{\varepsilon^{\max}}{m_{\mathrm{R}}}\frac{\mathrm{d}\phi_{\mathrm{R}}}{\mathrm{d}t}. \tag{S.13}$$

We can substitute from Equation S.12 and Equation S.2 to get

$$\frac{\mathrm{d}\lambda(t)}{\mathrm{d}t} = \lambda(t)\left(\lambda_{\mathrm{r}} - \lambda(t)\right) \tag{S.14}$$

where $\lambda_{\mathrm{r}}$ is the steady-state rich medium growth rate. Substituting in Equation S.11 gives us the lag time for nutrient upshift;

$$T_{\mathrm{lag,R}} = \frac{1}{\lambda_{\mathrm{r}}}\log\left(\frac{\lambda_{\mathrm{r}}}{\lambda_0}\right). \tag{S.15}$$

Here $\lambda_0$ is the effective initial growth rate immediately after the upshift. We assume that immediately after upshift,

1. The cell elongates with elongation rate of the rich nutrient, i.e., $\varepsilon(\lambda) = \varepsilon(\lambda_\mathrm{r})$.
2. The fraction of active ribosomes is equal to that of steady state growth in the rich nutrient, i.e., $\phi_\mathrm{R}^\mathrm{act}(\lambda) = \phi_\mathrm{R}^\mathrm{act}(\lambda_\mathrm{r})$. This assumption holds since ribosomes can be activated on a much faster timescale than they are produced ([2]).
3. R sector is the size it was during growth on poor nutrient, i.e., $\phi_\mathrm{R}(0) = \phi_\mathrm{R,p}$.

Putting the above assumptions together lets us write the equation for $\lambda_0$:

$$\lambda_0 = \frac{\varepsilon(\lambda_\mathrm{r})}{m_\mathrm{R}} \phi_\mathrm{R,p} f_\mathrm{R,r}^\mathrm{act}, \tag{S.16}$$

where $f_\mathrm{R,r}^\mathrm{act} = \phi_\mathrm{R,r}^\mathrm{act}/\phi_\mathrm{R,r}$. The growth-rate dependent terms are $\varepsilon(\lambda_\mathrm{r})$ and $f_\mathrm{R}^\mathrm{act}(\lambda)$ . Zhu et al. [7] derived the dependence on growth rate explicitly:

$$\varepsilon(\lambda) = \varepsilon^\mathrm{max} \frac{\phi_\mathrm{R}(\lambda)}{\phi_\mathrm{R}(\lambda) + \phi_\mathrm{R}^\mathrm{0,max}} \tag{S.17}$$

and

$$f_\mathrm{R}^\mathrm{act}(\lambda) = 1 - \left( \frac{\phi_\mathrm{R}^\mathrm{0,max}}{\phi_\mathrm{R}(\lambda)} \right)^2 . \tag{S.18}$$

Substituting back in Equation S.16 we get

$$\lambda_0 = \frac{\varepsilon^\mathrm{max}}{m_\mathrm{R}} \frac{\phi_\mathrm{R,r}}{\phi_\mathrm{R,r} + \phi_\mathrm{R}^\mathrm{0,max}} \phi_\mathrm{R,p} \left( 1 - \frac{\left(\phi_\mathrm{R}^\mathrm{0,max}\right)^2}{\phi_\mathrm{R,r}^2} \right) . \tag{S.19}$$

Note that $m_\mathrm{R}$ is a constant, $\varepsilon^\mathrm{max}$ and $\phi_\mathrm{R}^\mathrm{0,max}$ are physiological parameters, and $\phi_\mathrm{R,r}$ and $\phi_\mathrm{R,p}$ can be determined from Equation S.4. For downshifts we assume that the synthesis of required metabolic enzymes determines growth rates. We use an analogous calculation for the kinetics as we do for upshifts:

$$\begin{aligned} \frac{\mathrm{d}\phi_\mathrm{P}(t)}{\mathrm{d}t} &= \frac{\kappa_\mathrm{n,p}}{\rho} \phi_\mathrm{P}(t) \left( \phi_\mathrm{P,p} - \phi_\mathrm{P}(t) \right) \\ &= \lambda(t) \left( \phi_\mathrm{P,p} - \phi_\mathrm{P}(t) \right) \end{aligned} \tag{S.20}$$

where we use Equation S.1 for the second equality. Note that in poor nutrient growth, we don't have a pre-allocation of metabolic resources, i.e., $f_\mathrm{p} = 1$. Differentiating Equation S.1 gives us

$$\frac{\mathrm{d}\lambda(t)}{\mathrm{d}t} = \frac{\kappa_\mathrm{n,p}}{\rho} \frac{\mathrm{d}\phi_\mathrm{P}(t)}{\mathrm{d}t} \tag{S.21}$$

and substituting from Equation S.20 gives us

$$\frac{\mathrm{d}\lambda(t)}{\mathrm{d}t} = \lambda(t) \left( \lambda_\mathrm{p} - \lambda(t) \right) . \tag{S.22}$$

Substituting in Equation S.11 and integrating gives us

$$T_\mathrm{lag,P} = \frac{1}{\lambda_\mathrm{p}} \log \left( \frac{\lambda_\mathrm{p}}{\lambda_0} \right) . \tag{S.23}$$

To determine $\lambda_0$ we assume that immediately after downshift,

1. The P sector is the size it was during steady-state rich nutrient growth, i.e., $\phi_\mathrm{P} = \phi_\mathrm{P,r}$.
2. The influx of nutrients is given by the fraction of pre-allocated metabolic resources for the poor nutrient during rich medium growth, i.e., $(1 - f_\mathrm{r})$.

This gives us

$$\lambda_0 = (1 - f_\mathrm{r}) \phi_\mathrm{P,r} \frac{\kappa_\mathrm{n,p}}{\rho} . \tag{S.24}$$

Note that $\rho$ is a constant, $\kappa_\mathrm{n,p}$ is an environmental parameter, $f_\mathrm{r}$ is a physiological parameter, and $\phi_\mathrm{P,r}$ is given by Equation S.5.

**Quantifying fitness**

Consider batch dynamics where communities grow for a fixed time $T$, chosen to be large enough such that nutrients are completely depleted, following which the culture is diluted and switched to a new nutrient. We consider alternating rich and poor nutrients of concentrations $c_{\mathrm{r}}$ and $c_{\mathrm{p}}$ respectively. The dilution factor is denoted by $D$. Consider a single culture growing on the poor nutrient, with biomass $M$ and a lag time $T_{\mathrm{p}}$. The nutrient is fully depleted at time $\mathrm{t_p}$. Then growth dynamics follow

$$M(t) = M(0)e^{\lambda_{\mathrm{p}}(t-T_{\mathrm{p}})} \tag{S.25}$$

for $T_{\mathrm{p}} \leq t \leq \mathrm{t_p}$. We assume the following dynamics for nutrient concentration:

$$\frac{\mathrm{d}c}{\mathrm{d}t} = -\frac{\mathrm{d}M}{\mathrm{d}t}. \tag{S.26}$$

Substituting we get

$$\frac{\mathrm{d}c}{\mathrm{d}t} = -M(0)\lambda_{\mathrm{p}}e^{\lambda_{\mathrm{p}}(t-T_{\mathrm{p}})}1_{[T_{\mathrm{p}},\mathrm{t_p}]}, \tag{S.27}$$

and we can solve for the depletion time:

$$\int_0^{c_{\mathrm{p}}} \mathrm{d}c = \int_{T_{\mathrm{p}}}^{\mathrm{t_p}} M(0)\lambda_{\mathrm{p}}e^{\lambda_{\mathrm{p}}(t-T_{\mathrm{p}})}\,\mathrm{d}t\,. \tag{S.28}$$

After integrating and some algebra we get the condition

$$\mathrm{t_p} = T_{\mathrm{p}} + \frac{1}{\lambda_{\mathrm{p}}}\log\left(\frac{c_{\mathrm{p}} + M(0)}{M(0)}\right), \tag{S.29}$$

and equivalently for the rich medium we have

$$\mathrm{t_r} = T_{\mathrm{r}} + \frac{1}{\lambda_{\mathrm{r}}}\log\left(\frac{c_{\mathrm{r}} + M(0)}{M(0)}\right). \tag{S.30}$$

Now we want to calculate the steady state value of $M(0)$: for this, after completing one poor-rich cycle, $M(0)$ must be constant. Let $M_{\mathrm{p}}(t)$ and $M_{\mathrm{r}}(t)$ denote the biomass of the population when growing on poor and rich nutrients respectively. Assume a cycle starts with poor nutrient, then we have after the poor-nutrient cycle,

$$DM_{\mathrm{r}}(0) = M_{\mathrm{p}}(\mathrm{t_p}) = M_{\mathrm{p}}(0) + c_{\mathrm{p}}. \tag{S.31}$$

After dilution and growing in the rich nutrient,

$$M_{\mathrm{r}}(\mathrm{t_r}) = \frac{M_{\mathrm{p}}(0) + c_{\mathrm{p}}}{D} + c_{\mathrm{r}}. \tag{S.32}$$

For steady state we have the equality

$$\left(\frac{M_{\mathrm{p}}(0) + c_{\mathrm{p}}}{D} + c_{\mathrm{r}}\right)\frac{1}{D} = M_{\mathrm{p}}(0), \tag{S.33}$$

giving us

$$M_{\mathrm{p}}(0) = \frac{c_{\mathrm{p}} + Dc_{\mathrm{r}}}{D^2 - 1}. \tag{S.34}$$

We can substitute this in Equation S.31 to get

$$M_{\mathrm{r}}(0) = \frac{c_{\mathrm{r}} + Dc_{\mathrm{p}}}{D^2 - 1} \tag{S.35}$$

as the steady-state initial mass for rich medium. We can now substitute these in Equation S.29 and Equation S.30 to get the depletion times in poor and rich nutrients,

$$\mathrm{t_p} = T_{\mathrm{p}} + \frac{1}{\lambda_{\mathrm{p}}}\log\left(\frac{D(c_{\mathrm{r}} + Dc_{\mathrm{p}})}{c_{\mathrm{p}} + Dc_{\mathrm{r}}}\right) \tag{S.36}$$

and

$$\mathrm{t_r} = T_\mathrm{r} + \frac{1}{\lambda_\mathrm{r}} \log\left(\frac{D(c_\mathrm{p} + Dc_\mathrm{r})}{c_\mathrm{r} + Dc_\mathrm{p}}\right). \tag{S.37}$$

Consider an invading strain in this setup. Denote all parameters of the invader with a tilde; e.g., $\tilde{\lambda}, \tilde{M}$. When the resident population is at steady-state, the depletion time in either nutrient is driven entirely by the resident, given by Equation S.36 and Equation S.37. Denote the cycle number using a superscript. Then, post-dilution immediately after one rich-poor cycle,

$$\tilde{M}_\mathrm{p}^2(0) = \frac{\tilde{M}_\mathrm{p}^1(0)}{D^2} e^{\tilde{\lambda}_\mathrm{p}\left(\mathrm{t_p} - \tilde{T}_\mathrm{p}\right)} e^{\tilde{\lambda}_\mathrm{r}\left(\mathrm{t_r} - \tilde{T}_\mathrm{r}\right)}. \tag{S.38}$$

For invasion, the proportion of the invading strain must increase, giving

$$\frac{\tilde{M}_\mathrm{p}^2(0)/M_\mathrm{p}^2(0)}{\tilde{M}_\mathrm{p}^1(0)/M_\mathrm{p}^1(0)} > 1. \tag{S.39}$$

Since the resident is at steady state, $M_\mathrm{p}^2(0)/M_\mathrm{p}^1(0) = 1$ and the condition for invasibility reduces to

$$\frac{\tilde{M}_\mathrm{p}^1(0)}{D^2 \tilde{M}_\mathrm{p}^1(0)} e^{\tilde{\lambda}_\mathrm{p}\left(\mathrm{t_p} - \tilde{T}_\mathrm{p}\right)} e^{\tilde{\lambda}_\mathrm{r}\left(\mathrm{t_r} - \tilde{T}_\mathrm{r}\right)} > 1 \tag{S.40}$$

or

$$\tilde{\lambda}_\mathrm{p}\left(\mathrm{t_p} - \tilde{T}_\mathrm{p}\right) + \tilde{\lambda}_\mathrm{r}\left(\mathrm{t_r} - \tilde{T}_\mathrm{r}\right) > 2\log D. \tag{S.41}$$

Table 1: Parameters used in simulations

| **Parameter** | **Description** | **Value** | **Units** |
|---|---|---|---|
| $\kappa_{\mathrm{n,p}}$ | Poor nutrient quality | 0.15 | $\mathrm{h}^{-1}$ |
| $\kappa_{\mathrm{n,r}}$ | Rich nutrient quality | 6 | $\mathrm{h}^{-1}$ |
| $\mathrm{c_p}$ | Poor nutrient concentration | 1 | arbitrary |
| $\mathrm{c_r}$ | Rich nutrient concentration | Environmental parameter | arbitrary |
| $D$ | Dilution factor | Environmental parameter | - |
| $\varepsilon^{\max}$ | Maximal elongation rate | 18.72 | a.a./s |
| $\phi^{\max}$ | Maximal ribosomal allocation | 0.5 | - |
| $\phi_\mathrm{R}^{0,\max}$ | Maximal inactive ribosomes | Allocation parameter | - |
| $f_\mathrm{p}$ | Poor nutrient metabolic allocation | 1 | - |
| $f_\mathrm{r}$ | Rich nutrient metabolic allocation | Allocation parameter | - |
| $\mathrm{m_R}$ | Mass of a ribosome | 12251 | a.a. |
| $\rho$ | Conversion factor | 0.76 | - |

#### Data for Figure 1a and Figure 5b

In order to demonstrate variability in growth rates of closely related strains, we chose datasets that grew the same set of strains on multiple nutrients and measured the steady-state growth rates. The relevant datasets are [1] for *E. coli* and [4] for *V. splendidus*. The actual growth rates plotted in the figures can be found in the file `fig1a_data.csv` in the dataset associated with this paper, at `https://github.com/arjun-projcode/Code-envhist`.
